# Habitat Selection by the Ringed Kingfisher (*Megaceryle torquata stictipennis*) in Basse-Terre, Guadeloupe: Possible Negative Association with Chlordecone Pollution

**DOI:** 10.1101/2020.04.15.038729

**Authors:** Pascal Villard, Alain Ferchal, Philippe Feldmann, Claudie Pavis, Christophe Bonenfant

## Abstract

In the Lesser Antilles the Ringed Kingfisher *Megaceryle torquata stictipennis* is found on Martinique, Dominica and Guadeloupe. On Martinique and Guadeloupe, farmers growing bananas have made massive use of chlordecone from 1973 until 1993. Chlordecone is a remnant organochlorine insecticide which bioaccumulates in organisms easily. We carried out a survey of the Ringed Kingfisher in 2009 on Basse-Terre, Guadeloupe, traveling 270 km on 44 rivers to assess the effects of chlordecone on its habitat selection behaviour. We recorded 47 encounters of Ringed Kingfishers over the survey. A habitat selection analysis revealed that the Ringed Kingfishers were located mainly in sections of rivers flowing through the ombrophilous forest, but absent on the ocean shoreline or in the lower parts of rivers. The Ringed Kingfisher’s distribution on Guadeloupe could be negatively associated with areas heavily polluted by the chlordecone. We propose that the widespread use of chlordecone in banana plantations on Guadeloupe between 1973 and 1993 has adverse ecological consequences, and may be responsible for the absence of Ringed Kingfishers in lowland habitat on the island.

**Résumé:** Dans les Petites Antilles, le Martin-pêcheur à ventre roux *Megaceryle torquata stictipennis* ne vit que sur trois îles: Martinique, Dominique et Guadeloupe. En 2009, nous avons effectué un recensement en Guadeloupe, sur 270 km parcourus dans 44 rivières. Douze territoires de Martin-pêcheur à ventre roux ont été trouvés, et la taille totale de la population est estimée à 54-64 individus. Une analyse de la sélection de l’habitat montre que le Martin-pêcheur à ventre roux se trouvait principalement sur les rivières dans la forêt ombrophile et non pas sur le bord de l’océan ni dans la partie basse des rivières. En Guadeloupe, le Martin -pêcheur à ventre roux est négativement associé avec les surfaces hautement polluées par un insecticide organochloré, la chlordécone. Nous suggérons que l’épandage massif de la chlordécone de 1973 à 1993, dans les bananeraies de Guadeloupe, doit être grandement responsable de l’absence du Martin-pêcheur à ventre roux dans les habitats de basses altitudes de l’île.

**Resumen:** En las Antillas Menores, el Martín Gijante Neotropical *Megaceryle torquata stictipennis* solo vive en tres islas: Martinica, Santo Domingo y Guadalupe. En 2001, realizamos un censo en Guadalupe sobre 270 km recorridos en 44 ríos. Doce territorios de Martín Gijante Neotropical han sido registrados, y el volumen total de la población se estima a 54-64 individuales. Un análisis de la selección del hábitat muestra que el Martín Gijante Neotropical se encontraba principalmente en los ríos en el bosque lluvioso y no a la orilla del mar tampoco en la parte baja de los ríos. En Guadalupe, el Martín Gijante Neotropical esta negátivamente asociado con las superficies muy poluadas por un insecticida organicloro, la chlordecona. Sugeremos que la difusión masiva de la chlordecona entre 1973 y 1993, en las plantaciones de plátanos de Guadalupe, debe de ser altamente responsable de la aucencia del Martín Gijante Neotropical en los hábitats de baja altitud en la isla.

---

The Ringed Kingfisher *Megaceryle torquata* is a quite common and widely distributed species across Central and South America (Woodall 2001). In the Antilles, however, the presence of the species is heterogeneous among islands. For instance, the Ringed Kingfisher does not occur in the Greater Antilles, and in the Lesser Antilles *M. t. stictipennis* is found on Dominica, Martinique and Guadeloupe (Remsen 1990), suggesting that favorable ecological conditions are met on a few islands only, or that anthropogenic activities could be limiting the Ringed Kingfisher distribution. Of particular interest for the distribution of many bird species are the agricultural practices known to affect many aspects of their biology and ecology (see Geiger *et al*. 2010 for a review). Banana production on Guadeloupe, an important and intensive agricultural activity of many people living on tropical islands, have among other things led to marked modifications of the landscape like the widespread deforestation.

Tightly associated to banana farming is chlordecone, an organochlorine insecticide, that has been used heavily and extensively by farmers to control regular damage by a root borer, the banana weevil *Cosmopolites sordidus*, from 1973 until its ban on Guadeloupe in September 1993 by French law (Cabidoche and Lesueur 2011). The widespread use of chlordecone resulted in extended pollution of soils, waters and riverbed sediments (Crabit *et al*. 2016), which ecological consequences remains poorly documented and studied on Guadeloupe. Since spreading of chemicals in the landscape for agriculture can potentially alter the distribution of birds (Douthwaite 1982; Parsons *et al*. 2010; Mineau and Whiteside 2013), one factor that may have influenced Ringed Kingfisher populations in the Guadeloupe could be the historical use of chlordecone.

Chlordecone is a long lasting organic pollutant with carcinogenic, mutagen and/or reprotoxic consequences for exposed organisms. On wildlife, negative effects of chlordecone were reported with strong effects on the reproductive biology of many species (see Eroschenko 1981 for a review). In birds, bioaccumulation of chlordecone manifested itself in the eggs of osprey *Pandion haliaetus* living along a chlordecone-polluted river in the USA (Wiemeyer *et al*. 1988). Besides, organochlorides can damage the nervous system altering the breeding behavior of seabirds (Schreiber and Burger 2002). Organochlorides such as chlordecone can affect negatively bird biology to a large extent, ranging from organism functioning, to reproductive success, scaling up to ecological processes such population abundance and viability (Saaristo *et al*. 2018). For instance the reproductive failure induce by DDT because of fragile egg shells was accounted for the decreased in population abundance of the brown pelican *Pelicanus occidentalis* California, USA (Risebrough *et al*. 1971).

Lying at the top of the aquatic food chain, bioaccumulation of chlordecone in body tissues should be particularly acute for Ringed Kingfishers (Clarkson 1995). Two types of ecological consequences of pesticides are usually recognized in the literature (Saaristo *et al*. 2018). The direct effects act on the physiology of exposed organisms to pollutant in turn altering the metabolism, the cognitive ability or the behavior of individuals, and ultimately leads to reduced survival and reproduction. Indirect effects are generated by cascading effects of pollution on the abundance of preys or by changes in the habitat quality, which indirectly impacts on reproduction and survival of individuals of higher levels in the food chain (Campbell *et al*. 1997). Indirect effects are expected to modify the behavior of individuals, such as lowered predator avoidance or predation efficiency, or decreased movement ability, which all contributes to magnify the direct effects of contaminant on populations (Saaristo *et al*. 2018). Given reported adverse effects on the life histories of birds of many organochlorine compounds, especially for fish-eating birds in general (Moore 1965; USEPA 1975; Schäfer *et al*. 2011), the historical use of the chlordecone could explain the distribution and habitat selection of Ringed Kingfisher on Guadeloupe. Accordingly, many kingfisher species are used as biomonitors of waterways polluted by various contaminants including mercury (White and Cristol 2014, Zamani *et al*. 2009) and organochlorides such as DDT (Tanabe *et al*. 1998, Evans and Bouwman 2000).

Here we investigated whether chlordecone pollution could be related to Ringed Kingfisher habitat selection. We conducted presence-absence surveys for this species on 44 rivers located on Basse-Terre island, Guadeloupe, and calculated selection ratios to determine whether Ringed Kingfishers avoid polluted areas. If chlordecone negatively affects the abundance of Ringed Kingfishers, they should avoid areas with chlordecone contamination, *i*.*e*. we should find proportionally less individuals in the polluted areas than its availability in the landscape (Manly *et al*. 2007). To lend further support to that the potential negative association between chlordecone pollution and Ringed Kingfisher habitat selection, we accounted for those ecological variables that we believe would capture the main spatial structure of Basse-Terre in terms of available habitats and river characteristics, and could potentially confound the effect of chlordecone..

## Methods

We carried out our study in Guadeloupe, an archipelago of the Lesser Antilles in the Caribbean sea. On Basse-Terre island (848 km^2^, 16°10’ N 61°40’ W), a mountain range runs from north to south mostly between 500-800 m above sea level. The highest point is the 1,467 m summit of the southern volcano La Soufrière. Because the mountain range blocks the prevailing easterly winds from the Atlantic ocean, the east coast of Basse-Terre is wetter with a drier leeward west coast. Two main forest types have been described along the elevation gradient of the volcano (Van Laere *et al*. 2016). At low elevation lies the lower rainforest dominated by *Amanoa caribaea, Tapura latifolia* and *Dacryodes excelsa* trees. At higher elevation, the rainforest is mostly dominated by *Richeria grandis*. Many rivers run from the mountain range to the ocean, especially along the wetter east coast (Fig. 1b). We sampled 21 rivers on the windward east coast and 23 rivers on the leeward west coast; with 14 rivers (< 5 km), 15 (5 <10 km), 13 (10 < 25 km) and the 2 longest rivers (> 25 km). For the shorter 42 rivers, looking on a map of Basse-Terre, we selected the rivers to be equally spread around the island and with different levels of chlordecone pollution (Fig. 1a).

**Fig. 1:**
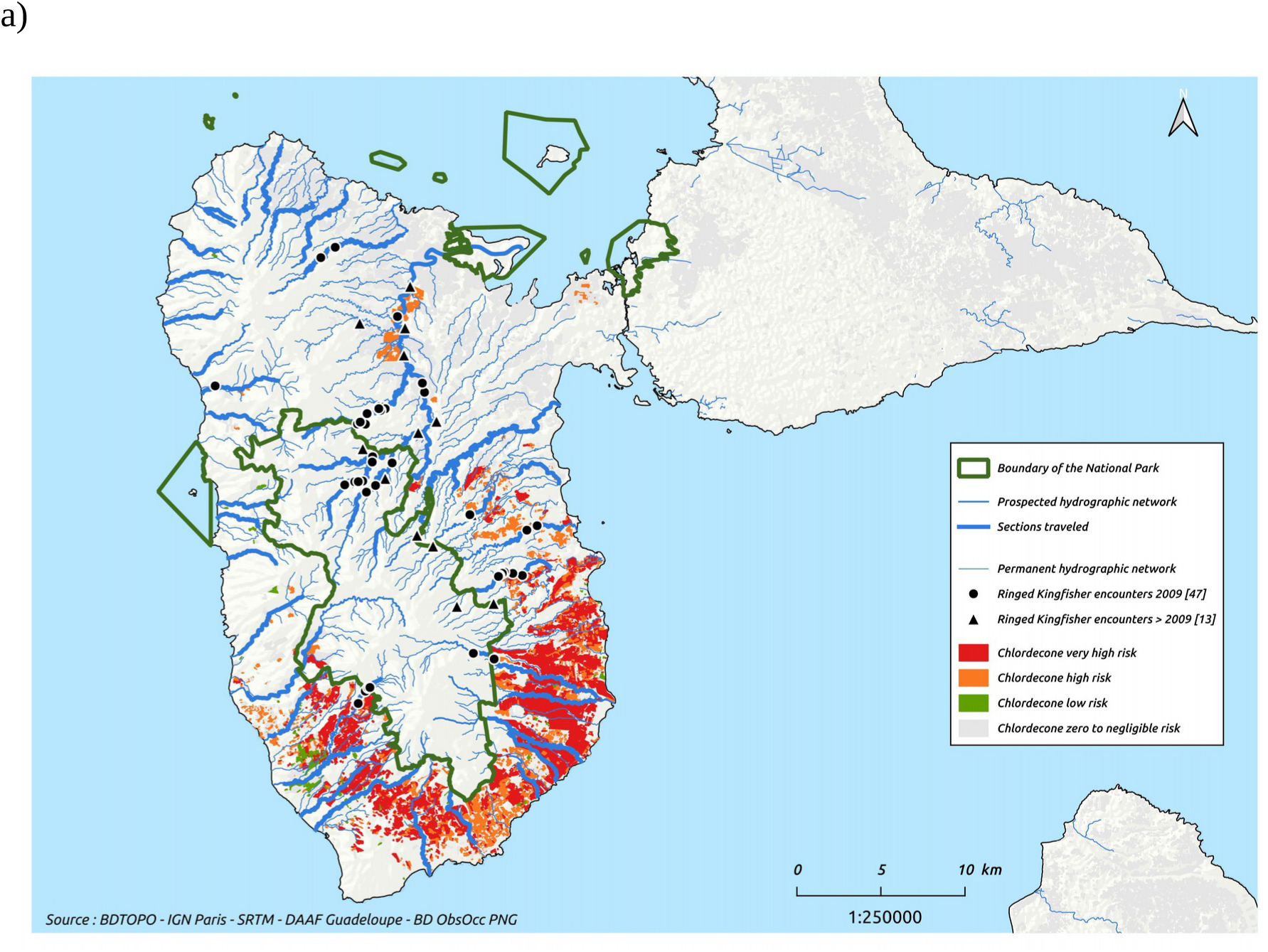
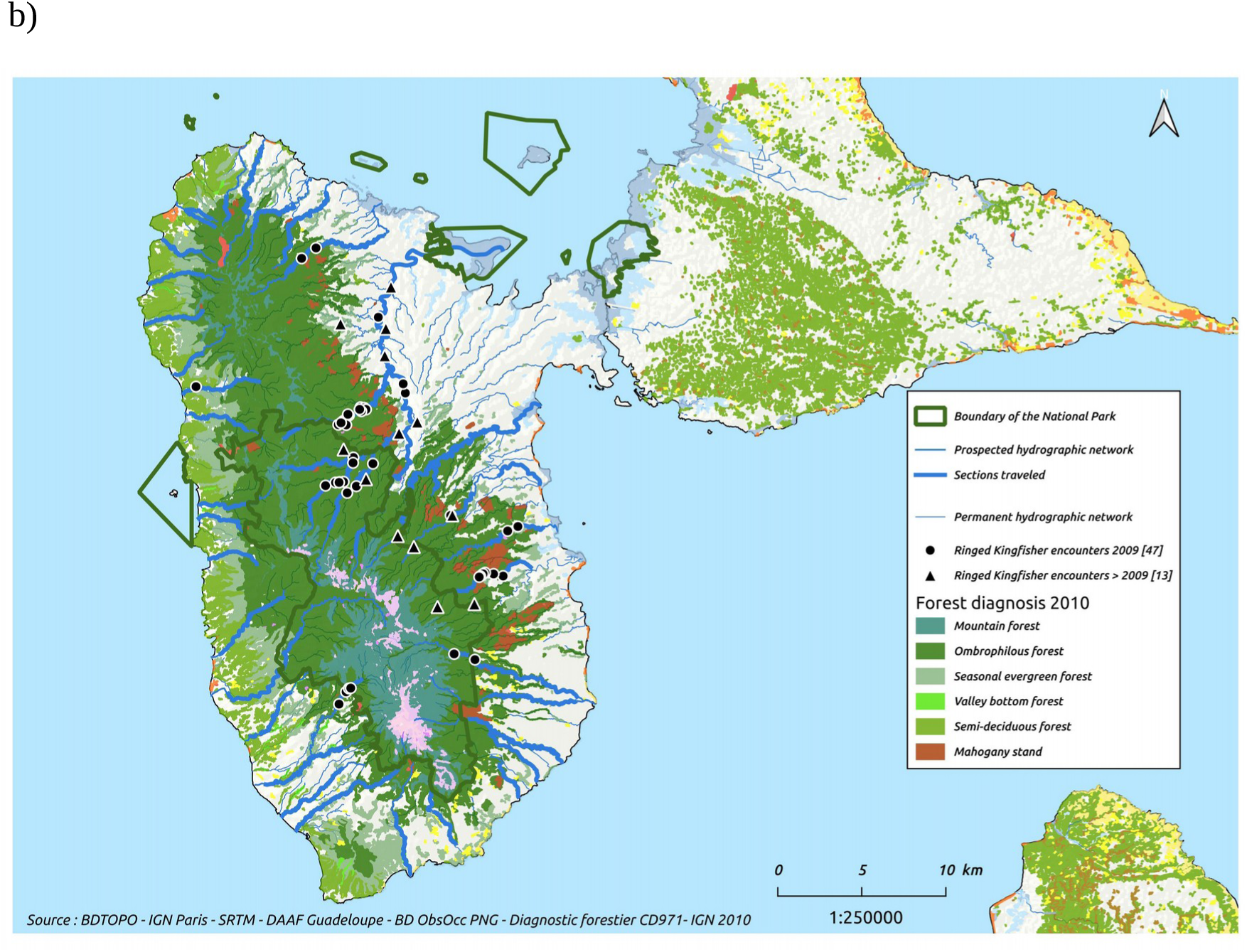
Location of Ringed Kingfisher (*Megaceryle torquata stictipennis*) encounters (2009 to 2017) on Basse-Terre, Guadeloupe (source for maps: BDTOPO – Institut Géographique National Paris; SRTM – DAAF Guadeloupe). In total, we recorded 47 observations of Ringed Kingfishers but the number of different birds remains uncertain, because some indivdiuals might have been seen several times if disturbed by the observer. A gross estimate of the number of different birds we have detected is 16. We overlappedthe observation locations on the map of (a) chlordecone exposure risk and (b) the map of forest habitats.

We conducted fieldwork in search of Ringed Kingfishers from 20 March to 20 August 2009. To obtain a baseline for the bird habitat use, we first studied the behavior of one pair, observing the birds for 33.5 hours over four days (on March 20^th^-23^rd^-24^th^ and 26^th^ 2009). We then surveyed Ringed Kingfisher using two methods: from kayak where possible in the lower sections of rivers or by walking along rivers. We assessed the Ringed Kingfisher presence on a river section by moving upstream slowly mostly in the river bed at a speed of 0.8 km per hour (1 km/h with a kayak), and starting at sunrise. We sampled the rivers only once and usually walked down away from rivers back to the coast afterwards. During the survey, we searched for the birds in the vegetation of riverbeds (with binoculars *Swarovski* 10×25) to spot any perched kingfisher. We recorded the GPS location of any Ringed Kingfisher heard or seen. Our experience showed that when a Ringed Kingfisher was present, by being careful enough (watching and listening) during the survey, we were quite confident not to miss the bird. Because some birds flew ahead of us as we surveyed rivers, we sometimes made repeated observations of a single individual. We always recorded the locations where Ringed Kingfishers were initially flushed. If the bird flew upstream, we also recorded the subsequent locations where it perched. We visually kept track of the flying bird until it reached its territory limit and turned back. We used this behavior to differentiate Ringed kingfishers individually, and any new bird later flushed upstream was considered as a different one. We acknowledge here that birds may have different habitat use between early spring and summer that could affect our results, even if it could be easier to see adult birds flying when feeding nestlings. Besides, because birdwatchers have been recording new, opportunistic Ringed Kingfishers observations between 2013 and December 2017 since our study was conducted (Karunati data base, accessed on May 10^th^ 2019). We retrieved the observation location reported by the observers from the Internet to reinforce our own observations and increase our sample size for the statistical analyses. We hence assumed observers spotted different Ringed Kingfishers, a likely met assumption given the rather large distance between the different observations we retained (mean: 7.1 km; range: 2.2 and 12.2 km). During the survey, we also made some anecdotal observations on adults Ringed Kingfisher bringing preys to their chicks and identified what species they feed on.

We described the local environmental conditions faced by Ringed Kingfisher along the sampled rivers and its surroundings with four ecological variables. With a GIS, we first created 250 m long river segments, starting from the river mouth toward its spring. For each segments we used a digital elevation model to calculate (*i*) the height difference in degree (*pente_moy*) between the start and the end of the river segment, which is associated with the speed, depth and turbidity of the waters and which we thus believe is associated with the fishing conditions for the birds; (*ii*) the average elevation in m of the 250 m long river segment (*alt_moy*). We believe this accounts for the elevation gradient in plant composition and land use by farmers and other human activities, because at high elevation the rainforest dominates, while in the lowland, forests were often cleared for banana and crops plantation, particularly on the east coast; (*iii*) the bearing in degrees (*angle*). We believe this accounts for the contrasting ecosystems between the windward and leeward coasts, because ombrophilous forest is generally found on both coasts, while seasonal evergreen and semi-deciduous forest, in contrast, are found only on the west coast (Van Laere *et al*. 2016). Finally, we added (*iv*) the chlordecone pollution level according to the classification and map of land pollution risks by Rochette *et al*. (2017). Rochette et al. (2017) assessed soil pollution using a multiresidue analysis of pesticides of 227 samples on randomly chosen rivers plus 120 samples collected at the estuary of 59 different watersheds, and interpolated this data with a Kriging procedure to produce a map of chlordecone pollution risks (Rochette *et al*. 2017). Class 5 soils have “very high risk” of chlodercone pollution (>90%) covering 4,761 ha (5.6% of Basse-Terre land); a class 4 “high risk” pollution (∼80%) making 1,629 ha (1.9%); class 3 soil has “low risk” of pollution (∼30%) covering 181 ha (0.2%); Class 2 soils have “low to negligible risk” of chlordecone pollution risks (<10%), covering 27,929 ha of the island (32.9%); Finally the last class 1 should show “no pollution by chlordecone”, representing the remaining 59.3% of the Basse-Terre area. We assessed the pollution risk of each river segment (*val_tc_chlor*) by assigning it the highest pollution risk class of all contiguous areas.

To explore whether Ringed Kingfishers avoided polluted areas, we calculated selection ratios (noted SR), a classical measure of preference and avoidance in habitat selection studies (Manly *et al*. 2007). An SR is calculated by dividing the proportional use of a given habitat by the proportional availability of that habitat. An SR value of one means that birds use the focal habitat randomly (i.e., use is equal to availability). When individuals avoid a particular habitat, it will be used to a lower extent than its availability and the SR will be <1. Conversely, a preferred habitat will have a SR >1. Note that habitat preference and avoidance are the standard terms used in habitat selection studies and describe an observed pattern that can be the result of many different behaviors and factors—e.g., the active avoidance of a threat, or the absence of a species in a certain habitat due to a lack of suitable resources. For all sampled river segments, we calculated the proportions falling into each of the five chlordecone pollution risk classes. We repeated this calculation for the subset of river segments where we observed Ringed Kingfishers. We then calculated selection ratios for each pollution class by dividing, for a given pollution class, the proportion of river segments with Ringed Kingfisher observations by the proportion of all river segments in that class. We computed the variance for each SR according to Aho and Bowyer (2015) for the variance of the two ratios.

We complemented the selection ratio analysis of Ringed Kingfisher with a multivariate approach called the MADIFA (Mahalanobis Distance Factorial Analysis, Calenge *et al*. 2008). A principal component analysis is first performed on the environmental descriptors to uncover the most influential variables structuring the landscape. This multivariate description of the environment corresponds to the ecological space available to the birds. Then the method compares, in the ecological space defined above, where birds are present and absent. If birds select the habitat along some variables, the locations where birds were detected should be associated with the one or several axes. An advantage of the method over the selection ratios (Manly *et al*. 2007) is that spatial covariation among the different environmental descriptors we included is visualized and accounted for by the MADIFA. As a side note, none of the methods could account for potential pseudo-replication in the data arising from the repeated observations of the same individuals. We hence treated each observation as independent record, and assumed little or no effect of pseudo-replication on our results. We ran all analyses using R 3.5 (R core team 2019) and the associated package *asbio* (Aho 2019) and *adehabitatHS* (Calenge 2006).

## Results

We surveyed 270 km of river habitat over a period of 376.6 hours (93.8% of the habitat by foot and 6.2% by kayak). Thus we surveyed 69% of the total length (395 km) of the 44 rivers on Basse-Terre that we sampled representing 27% of the total length of permanent rivers on Basse-Terre that were available to the Ringed Kingfisher. We recorded a total of *n* = 47 Ringed Kingfisher locations. From these observations, we tentatively assessed the possible number of different birds to 16 among which 12 were seen as singletons, and 2 couples.. On Basse-Terre the total length of rivers with permanent water was 1,012 km. During our survey we noticed that rivers shorter than 2 km long and representing 113 km out of the 1,012 km were not occupied by Ringed Kingfishers. Thus, river habitat available to Ringed Kingfishers that we later used for our habitat selection analyses reduced to 899 km long (including 395 km for the 44 rivers sampled). Since our study was conducted, 13 new Ringed Kingfisher observations have been recorded between 2013 and 2017 by birdwatchers (Fig. 1a). Note that these new locations where birds were seen fit in our Ringed Kingfisher habitat selection patterns of an avoidance of polluted river segments whether these observations were made in areas we previously visited during our sampling (9 observations) or not (4 observations)..

We overlaid the data on Ringed Kingfisher distribution with the map of level of chlordecone contamination risk (Fig. 1a) and forest habitat (Fig. 1b). Our results suggest that Ringed Kingfishers (*i*) are found outside of high chlordecone contamination areas (> 80% polluted, classes 4 and 5 of chlordecone contamination level), and (*ii*) mostly in the ombrophilous forest. The statistical analysis of habitat selection confirmed the visual inspection of Ringed Kingfisher locations on the map. River segments with no chlordecone were used slightly more relative to availability (SR_1_ = 1.18 ± 0.01; Fig. 2). Conversely, Ringed Kingfisher used river segments with any risk of chlordecone pollution significantly less relative to its availability (for all river segments with pollution risks of 2 to 5: SR_2–5_ = 0.46 ± 0.30, p = 0.07). Biologically speaking it means that the odds to find a Ringed Kingfisher are 2.56 higher in unpolluted than in polluted river segments on Basse-Terre, on average. When considering each pollution level separately, all estimates of selection ratios of polluted river segments were < 1, although not statistically different from one (see Fig 2; SR_2_= 0.32 ± 0.98; SR_3_ = 0.00 ± 2.15; SR_4_ = 0.00 ± 2.08; SR_5_ = 0.90 ± 0.50).

**Fig. 2:**
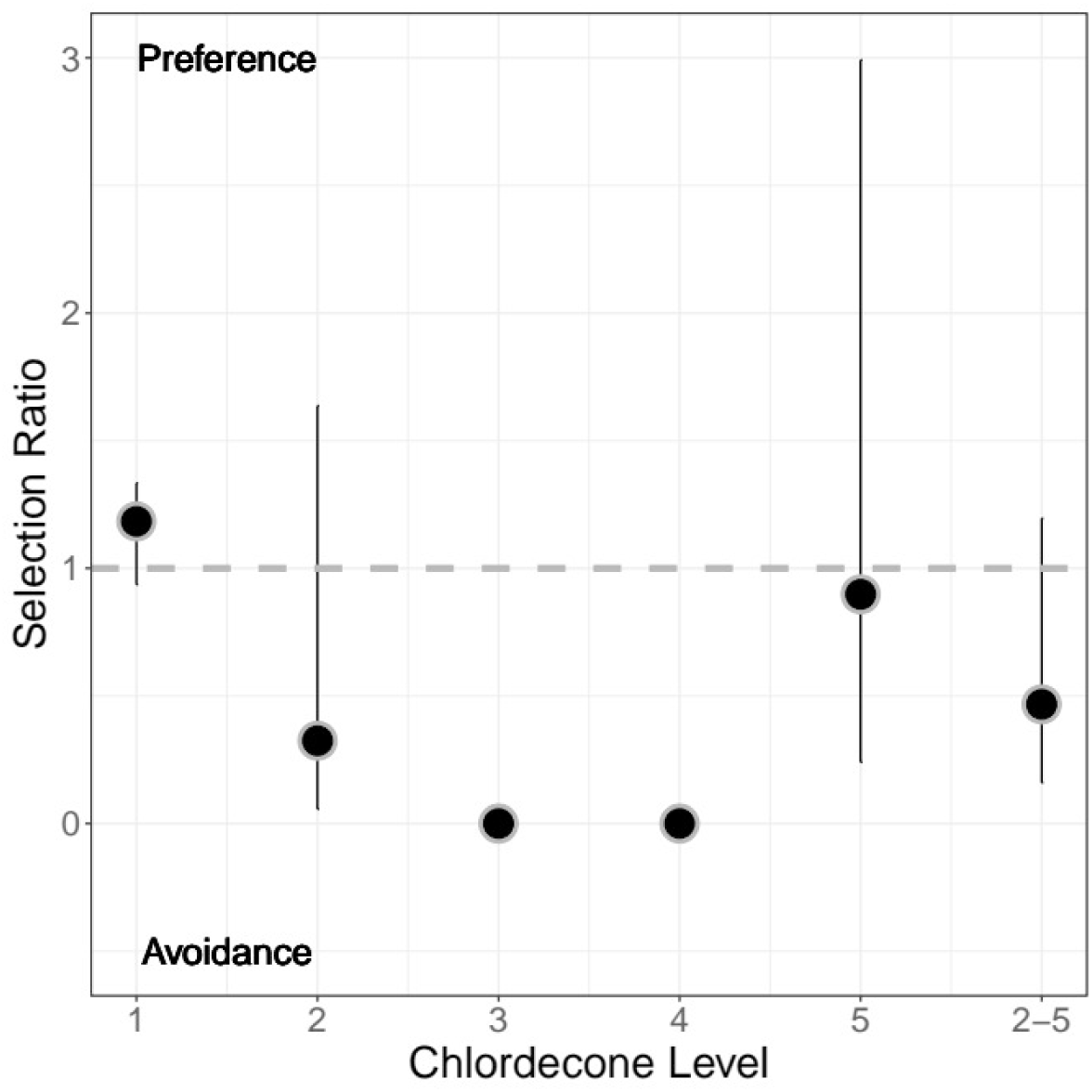
Selection ratios (with associated 95% confidence limits) based on Ringed Kingfisher observations made along river segments with five different levels of chlordecone pollution on Basse-Terre, Guadeloupe between 2009 and 2017 (*n* = 47 bird locations on approximately 16 different individuals). Chlordecone levels from 1 to 5 are increasing risks to find polluted river segments, while the last category pooled all polluted segments whatever its risks. Although selection ratios are not statistically different from 1, Ringed Kingfishers seem to use non-polluted segments of rivers (chlordecone level 1) more relative to availability, contrasting with river segments with low to high levels of pollution (chlordecone level 2 to 5) which are used less relative to availability.

The multivariate habitat selection analyses lend further supports to the selection ratio results. The correlation circle (Fig. 3A) showed that elevation and bearing correlated positively because high elevation areas are found mostly in the southwest of the island. Importantly, the level of chlordecone pollution was orthogonal to the other environmental variables, suggesting that the use of the pollutant in the landscape has been (statistically) independent of the other landscape variables used to described the Ringed Kingfisher environment. It also means that any effect of chlordecone we would find on the Ringed Kingfisher habitat selection would not be the outcome of potential confounding effect of the described landscape structure. When projecting the points on the first two axes of the ecological space, the MADIFA shows that Ringed Kingfisher (black dots, Fig. 3B) selected high elevation river segments rather than low elevation segments. Similarly, river segments with too high a denivelation were used less intensively than flatter ones. Furthermore, none of the river segments where birds were seen are on the lower-left quarter of the plot, which represents the ecological space with chlordecone pollution (black dots, Fig. 3B)..

**Fig. 3:**
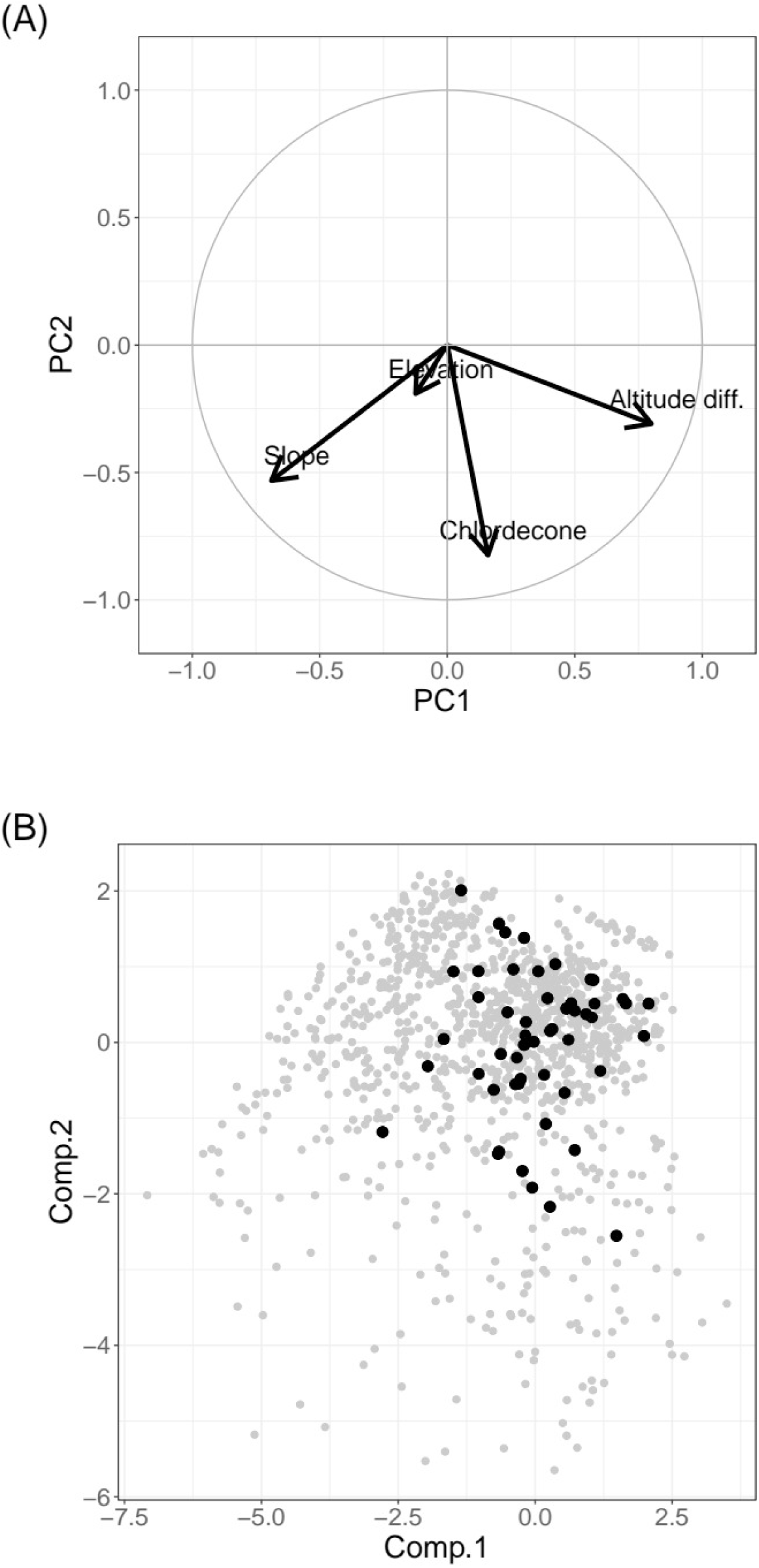
Graphical output of the MADIFA (Mahalanobis Distance Factorial Analysis, Calenge *et al*. 2008) displaying the ecological conditions prevailing in river segments where Ringed Kingfishers (*Megaceryle torquata stictipennis*) were observed (black dots) compared to available river segments on the island (grey dots), on Basse-Terre, Guadeloupe, between 2009 and 2017 (*n* = 47 bird locations on approximately 16 different individuals). We used 4 variables to describe the environmental condition for each 250 long river segments: the average height difference in degree (Elevation diff.), the average elevation in m (Elevation), the bearing in degrees (angle), and the chlordecone pollution level (chlordecone). (a) Correlation circle of the principal component analysis of the river segments characteristics showing that chlordecone pollution is only weakly associated with other environmental descriptors we used; (b) the MADIFA shows that none of the river segments where birds were seen were found on the upper-right quarter of the plot (black dots), which represents the ecological space with low to high chlordecone pollution.

Nonetheless, coupled with the selection ratios, this multivariate analysis is congruent with the idea that on Guadeloupe, the Ringed Kingfisher might use river segments polluted by chlordecone less than their availability. Accordingly, none of the river segments where birds were seen were found on the upper-right quarter of the plot (black dots, Fig. 3B), which represents the ecological space with chlordecone pollution.

## Discussion

Given the particular agricultural history of some islands in the Antilles, we put forward the hypothesis that long-lasting effects of chloredone pollution, which persists in the environment for centuries (Crabit *et al*. 2016), could have had adverse ecological conditions on freshwater ecosystems of Guadeloupe, including the Ringed Kingfisher. On Basse-Terre, we document a lower relative use of the lower sections of rivers by the Ringed Kingfisher, which run though the heavily chlordecone contaminated estuary on the east coast (Fig. 1). A study carried out on the European Dipper (*Cinclus cinclus*) found a similar absence of this river-dwelling species in 93.7% of polluted or strongly polluted streams in Italy (Sorace *et al*. 2002). Chlordecone pollution could therefore at least partly explain the past and current limited distribution of Ringed Kingfishers on Basse-Terre that we describe here (Fig. 1a). In the 1970’s, Ringed Kingfishers were regularly recorded in Sofaïa, Capesterre and Trois-Rivières on this island (Pinchon 1976). Forty years later, we detected the species in the first two locations still, but only in upstream areas; one possible explanation is that chlordecone has not been spread by banana farmers in upstream areas. Overall, the Ringed Kingfisher is not found in the lower part of rivers or the ocean front on Basse-Terre (Fig. 1). This could be related to the 84.1 % of watersheds analyzed having pesticides such as chlordecone (Rochette *et al*. 2017). Finally during the 10 years following our survey, birdwatchers recorded several sightings of Ringed Kingfishers on Guadeloupe, but these were all made outside of areas of high chlordecone pollution risks. Regarding the four Ringed Kingfishers detected in polluted area (class 4 and 5), they were found where land pollution was patchily distributed, with access to some unpolluted areas as well.

The spatial avoidance of polluted river segments by Ringed Kingfisher could result from direct and indirect effects of chlordecone on the birds. Direct effects usually lead to the death of individuals or regular reproduction failures because of organism dysfunction when exposed to chemical pollutants (Saaristo *et al*. 2018). Although anecdotal, we could observe the feeding of Ringed Kingfisher chicks with basket shrimp *Atya innocous*, stream shrimp *Macrobrachium crenulatum*, and the river goby *Awaous banana*. An ecolotoxicology study carried out on southern Basse-Terre river evidenced substantial amount of chlordecone in those three prey species (Coat *et al*. 2011). Although we currently lack of published material on the Ringed Kingfisher diet in the tropics to strengthen this observation, we can expect a direct effect of chlordecone *via* bioaccumulation in its tissue from polluted food resource in lowlands of Guadeloupe. Clearly, an epidemiology study on the Ringed Kingfisher is most needed to assert the physiological effects of chlordecone on the bird organism and life histories. Moreover, indirect effects of chlordecone may proceed from cascading effects of contamination on prey species community or size structure of the populations and, in turn, can lower the suitability of habitat for predators in terms of food resources (Saaristo *et al*. 2018). With no bird banding we cannot assess whether kingfishers actually move from one river to another to actively avoid chlordecone pollution, but over time both direct and indirect effects could have led to the absence of individuals from polluted areas as we suggest is the case on Guadeloupe.

Our results indicate that other ecological factors might also be related to the distribution of the Ringed Kingfisher on Guadeloupe. For example, river segments located on the eastern coast are used more than expected from its availability by Ringed Kingfishers (Fig. 3), and we observed the Ringed Kingfisher most on the windward east coast (Fig. 1b), where ombrophilous forests grow. Rivers running though this type of forest are long with gentle slopes, and river banks offer many clay cliffs to the birds for nesting. In this ecosystem, food resources are likely available to Ring Kingfishers given the many school fishes we saw while walking the rivers. The vegetation structure of the forest also provides numerous perches ideal for birds to fish. This ombrophilous forest hence appears a rather suitable habitat for the Ringed Kingfisher both from a food resource viewpoint, and for nest excavation too (Woodall 2001). Conversely, on the dryer leeward west coast fish abundance is lower and rocks make most of the river banks. On average rivers are also shorter (1.8 km *vs*. 3.0 km) and steeper (16.6° *vs*. 11.0°) hence presenting more rapid water streams. We noted previously that no kingfisher were using streams of < 2km long, maybe because it is too shallow for diving. Generally speaking the west coast habitat seems less favorable to Ringed kingfishers because of river characteristics, low food resource and the lack of soft banks forcing birds to fly far from waters to dig its burrow nest (Woodall 2001). Accordingly, we detected only two individuals on the west coast. One exception was an individual living in the northwest (Fig. 2), that we could easily associate with the presence of a fish farm where it is seen fishing regularly.

Elevation is another influential ecological variable of Ringed Kingfisher habitat selection. Here, elevation likely serves as a proxy of river physical characteristics, as water speed, depth, temperature or turbidity all vary gradually from the high elevation river source to the estuary (Allan 1995). It is known that the species needs fairly deep water to dive and should hence be most abundant in the lowlands (Skutch 1972). We found the opposite pattern and attributed this difference to a combination of chlordecone pollution and unsuitable habitats for Ringed Kingfishers. On Basse-Terre, although chlordecone pollution of soils and waters decreases with altitude (Fig. 1), our river sampling resulted in a weak spatial correlation between the two variables at the landscape level (Fig. 3a). Statistically speaking, we could decipher the effect of chlordecone pollution risks and of elevation, and found that Ringer Kingfisher was present relatively less in polluted river segments or in those located at low elevation (Fig. 3b). That the birds avoid unpolluted lowland river segments could be explained by the marked human footprints on the beds, banks and surroundings of rivers in Guadeloupe. For instance, the riverbed of the river flowing through the city of Basse-Terre is made of concrete mostly, which probably reduces fish abundance drastically. Similarly in banana plantations the forest is sometimes clear cut down to the river edge, except from a narrow strip of trees along river banks, here again modifying the vegetation structure and density (note that here a lack of perches could not explain the low relative use of Ringed Kingfishers in banana plantation). Lowlands of Guadeloupe islands are at the heart of human activities, which generates all kinds of perturbations for wildlife and possibly for the Ringed Kingfisher. Light pollution, chlordecone upstream pollution, active disturbance, presence of predators or competition for food with fishermen could all contribute to deter Ringed Kingfishers from lowland rivers beside the polluted river segments.

Our study suggests that the Ringed Kingfisher could be a useful bioindicator of areas with organochlorine pollution, and most specifically chlordecone. We contend that our habitat selection analysis lacks statistical power but given the potentially dramatic consequences for the Ringed Kingfisher, the low level of statistical significance should not hamper the biological signal we document here (see Yoccoz 1991 for a discussion of statistical significance). Although we cannot prove the generalized ecological effects of chlordecone in the Antilles, we advance that historical chlordecone use by banana farmers could have had and be still having an ongoing influence on Ringed Kingfisher populations and distribution on certain islands, including Guadeloupe. To further explore this potential issue, researchers should collect data on the ecology of the Ringed Kingfisher with a detailed descriptions of diet through the year, habitat, territory size, and behavior, and also to investigate the ecological consequences of chlordecone pollution could have on individuals, populations and avian communities.

## Acknowledgements

This work was supported by the national park of Guadeloupe, DEAL Guadeloupe (the Regional Department of Environment) and AEVA. We thank Stéphane Di Mauro for preys determination, Gavin Hunt for editing the English and an anonymous reviewer for his constructive comments. We thank Shirley Dolo for the Spanish version of the summary. We acknowledge the insightful and extensive help of Alice McBride who greatly improved our manuscript.

## Contributions

P.V. designed the study, carried out fieldwork and wrote the paper; A.F. made all maps and extracted with a SIG the data for analysis; P.F. and C.P. helped shape the report and C.B. performed the statistical analysis, wrote their interpretation and contributed to the writing.

